# APR-246 & COTI-2 increase chemoradiotherapy sensitivity via ROS-induced DNA damage and ferroptosis in p53 mutant HPV negative head and neck squamous cell carcinoma

**DOI:** 10.1101/2024.12.23.630069

**Authors:** Tycho de Bakker, Morgane Cogels, Chrysanthi Iliadi, Sebastien Penninckx, Philippe Martinive, Dirk Van Gestel

**Affiliations:** Jules Bordet Institute

**Keywords:** APR-246, COTI-2, Chemoradiotherapy, reactive oxygen species, Ferroptosis, DNA damage

## Abstract

About 80% of patients with HPV negative head and neck squamous cell carcinoma are diagnosed at stage III or IV. In these cases, Surgical resection followed by (chemo-)radiotherapy is the standard treatment for most cases. However, many patients present with unresectable and/or resistant disease and even metastases, limiting therapeutic options. A common molecular defect in these tumours is the deactivation of the tumour suppressor gene TP53, frequently through inactivating mutations. Restoring TP53 function, combined with the standard chemoradiotherapy may enhance the therapeutic outcomes.

In this study, the efficacy of two TP53 reactivating compound, APR-246 and COTI-2, was evaluated in combination with chemoradiotherapy on two different human HNSCC cell lines. Results highlight a synergistic effect of the combination treatment, significantly reducing clonogenic survival, spheroid growth and subcutaneous tumour growth in a preclinical murine model. Mechanistic investigation suggests that this effect is linked to redox imbalance caused by the generation of reactive oxygen species. This appears to play a key role in the Fenton reaction, further facilitated by an increase in DMT1 or decrease in FTH1 expression, leading to elevated cytosolic iron and lipid peroxide levels. Additionally, the reactive oxygen species may contribute towards the increase in both single and double strands breaks observed in several western blots. Overall, these results suggest that combining TP53 reactivation with chemoradiotherapy could trigger ferroptosis, improving tumour control in HNSCC.

## Introduction

In 2022, 660.000 people worldwide were diagnosed with head and neck squamous cell carcinoma (HNSCC), which accounted for about 4% of all cancer cases ^1^. These patients are classified by HPV status, with five-year overall survival rates differing significantly between HPV-positive patients (20- 40%) and HPV-negative ones (60-80%)^2^. Around 40 to 60% of HPV-negative patients are diagnosed at an advanced stage (III or IV)^3^. For these patients, the standard treatment consists of surgical resection followed by adjuvant (chemo-) radiotherapy (CRT)^4^. However, many patients experience resistance to the treatment or develop severe acute and chronic toxicities, such as mucositis, odynophagia & dysphagia and xerostomia, laryngeal dysfunction & ototoxicity ^5^. These adverse events result in reduction or discontinuation of chemotherapy or in a dose-de-escalation in radiotherapy^6^. Interestingly, about 70% of HPV-negative HNSCC patients present with somatic TP53 mutations, which often target the DNA binding domain rendering p53 non-functional^7^. Compounds such as Eprenetapopt (APR-246) and COTI-2 have been shown to reactivate certain mutated forms of p53 by inducing conformational changes in the altered DNA-binding domain, thereby restoring wild-type protein function ^8^. 2,2-bis(hydroxymethyl)-1-azabicyclo[2,2,2]octan-3-one was originally discovered by Bykov et al. who screened a library of small, low molecular weight molecules for their ability to suppress growth of p53 mutated cells ^9^. Named ”p53 reactivation and induction of massive apoptosis”, (PRIMA-1), the compound was later modified in 2005 to create a methylated derivative, PRIMA-1^Met^ (APR-246), which enhances cellular uptake by improving membrane permeability. Once inside the cell, APR-246 is converted into methylene quinuclidinone (MQ), a reactive electrophile able to covalently bind to cysteine residues 124 and 277 in the DNA binding domain of p53. This induces conformational changes that stabilizes the wildtype p53 conformation^10^. Notably, APR-246 also affects the redox balance by interacting with glutathione (GSH) as well as thioredoxin reductase 1 driven by its sulfhydryl binding properties^11^. It has been investigated in combination with various compounds across different cancer types. In HNSCC, re-sensitization towards conventional chemotherapeutics such as cisplatin^10^, 5-FU^12^, paclitaxel^13^ and erlotinib^14^ have been achieved. Additionally, agents such as PHEN^15^ (PARP inhibitor) and Piperlongumine^16^, which inhibits MAPK, NFkB and JAK/STAT signalling pathways, have shown to enhance effects of APR-246 in HNSCC.

In 2013, a thiosemicarbazone called COTI-2 was identified using the computer-aided drug design platform CHEMSAS. The molecule is effective at nanomolar concentration across different cancer types. In 2019, Lindemann et al. demonstrated that COTI-2 can reactivate p53 through its Zn^2+^ ion binding properties, which enable it to refold the DNA binding domain towards its wild type conformation, restoring its activity^17^. Upon its discovery, it was noted that COTI-2 also suppressed the growth of p53 wildtype cell lines, indicating p53 independent effects. These effects are attributed to the inherent properties of thiosemicarbazones being chelators. As such COTI-2 also binds copper ions, forming a Cu-COTI-2 complex that interacts with GSH and is subsequently exported by the ABCC1 transporters resulting in a redox imbalance ^18^.

Given the low 5-years survival rates and the frequent treatment resistance reported in advanced HNSCC patients, new therapeutic combinations that either enhance tumor control or reduce adverse events related to the treatment without compromising efficacy are needed. In this context, the present work investigate the effects of combining APR-246 and COTI-2 with standard chemoradiotherapy on HNSCC in *in vitro* and tumour-bearing nude mouse models.

## Materials & Methods

### Cell culture

Two p53 mutated HNSCC cell lines, FaDu (Y126-K132del & R248L), derived from a hypopharyngeal cancer, and Detroit-562 (R175H) derived from the pleural effusion of metastatic pharyngeal cancer ^19^ were purchased from American Type Culture Collection (ATCC). Both cell lines were cultured in DMEM supplemented with 5% FBS and 1% antibiotics (penicillin streptomycin and kanamycin), incubated at 37°C with 5% CO2. Cultured cells were regularly tested for Mycoplasma contamination (Mycoalert, Lonza, Switzerland) as well as submitted to an STR authentication (Eurofins, Germany).

### Reagents

APR-246, Buthionine sulfoximine (BSO), cisplatin, COTI-2 and erastin were purchased at Bioconnect, Netherlands. All treatment molecules used were dissolved according to the guidelines provided by the supplier. APR-246 was dissolved in PBS while BSO and cisplatin were dissolved in water. COTI-2 and erastin were dissolved in DMSO.

### *In vitro* irradiation

Cells were irradiated using the Elektra versa HD, with a 6MV photon beam at a dose rate of about 5.5Gy/min. For adequate backscattering absorption, well plates were placed on a 5 cm-thick polystyrene phantom. On top, a 6mm-polystyrene was placed to ensure dose homogenisation.

### MTT assay

MTT assays were conducted following the manufacturer’s instructions (Sigma Aldrich, Belgium). Three thousand cells were seeded in each well of a 96-well plate and incubated for 24 hours in 5% CO2 incubator at 37°C overnight. Cells were then treated with a range of concentrations of APR-246 (10^-7^- 10^-3^M), COTI-2 (10^-10^-10^-6^M) or cisplatin (10^-8^-10^-4^M). After 72 hours of treatment, the supernatant was removed, cells were washed with 100µl of PBS and incubated with 100µL of MTT (3-(4,5- Dimethylthiazol-2-yl)-2,5-Diphenyltetrazolium Bromide) for 2 hours. Removal of MTT and addition of 100µL of DMSO allowed release of the purple formazan which is subsequently measured using the Glomax® (Promega,Wisconsin) spectrophotometer at a wavelength of 560nm. Six technical replicates were seeded for each condition.

### Clonogenic assay

Clonogenic assay^20^ was performed by seeding 500,000 cells per well in a six well plate, allowing cells to attach for 24 hours before treating with APR-246 (100µmol/l), COTI-2 (50 nmol/l) and/or cisplatin (600nmol/l). The choice of each effector concentration was based on the IC20 values determined in the MTT assay and extrapolated for use in a six well plate. 24 hours later, each plate was irradiated with one fraction of 0, 2, 4 or 6 Gy. Subsequently, cells were harvested by trypsinization and reseeded for each dose condition at an appropriate cell density, 500 cells/well, 1,000 cells/well, 2,000 cells/well and 5,000 cells/well, respectively. After a 14 days incubation period, the colonies were stained with crystal violet in 2% ethanol. The amount of visible colonies (containing 50 or more cells) was considered to represent the surviving cells, which were counted manually. The plating efficiency (PE) and survival fraction were determined for each conditions using the following formula:

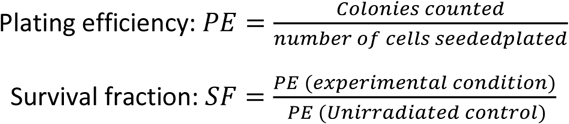

All clonogenic data were fitted using the linear-quadratic model:

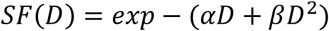

Where SF is the surviving fraction of the cells; α and β define the linear and the quadratic components, respectively, and D is the deposited dose. Sensitivity Enhancement Ratio (SER) was calculated at 20% and 50% survival as well as the coefficient of Drug Interaction for each conditions using the following equations:

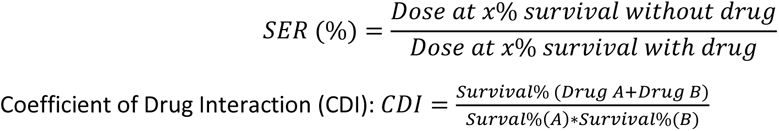

When CDI<1 the combination is considered synergistic, when CDI=1 the combination is considered additive and when CDI>1 combination is considered antagonistic.

### Tumour spheroid growth assay

The spheroid growth assay was performed as described by Friedrich et al.^21^. The spheroids were formed by seeding 5,000 GFP transfected cells in ultra-low attachment 96 well plates (Corning, New York). Subsequent centrifugation of the 96 well plate at 400g for 5 min allowed the cells to accumulate at the round bottom of the well. After 24 hours of incubation cells were exposed to APR-246 (60µM), COTI-2 (25nM) and Cisplatin (15 and 10µM for Detroit-562 and FaDu respectively), for five technical replicates in each condition. One day after treatment with the effectors, the plates were exposed to one fraction of 5 Gy irradiation. Afterwards, plates were placed in a live cell imaging machine, Incucyte S3® (Sartorius, Belgium) and data were registered for each well separately every 24 hours. Images obtained for both irradiated and control plates were analysed using the dedicated Incucyte® software to determine the GFP (+) area

### Annexin-FITC/7-AAD staining

50,000 cells/well were seeded in a six well plates. They were treated with APR-246 (100 or 50µM), COTI-2 (200nM) and Cisplatin (1µM). Additionally, half of the plates were irradiated with one fraction of 5Gy X-ray, while the other half of plates remained un irradiated. After 72 hours of incubation, cells were trypsinized and resuspended in 150µl of PBS. Finally, 5µl of Annexin V-FITC and 5µl of 7-AAD were added to each tube. After a 20 minute incubation, samples were tested on the Beckman Coulter Navios® flow cytometer (California, USA) until 10,000 events are measured or 10 minutes have elapsed.

### Western blot

Petri dishes were seeded with 1.5 million cells and treated with APR-246, COTI-2 or cisplatin either alone or in combination with a 5Gy radiation fraction. Depending on the protein to be tested, cells were lysed either 1, 16 or 24 hours after treatment. Bands were detected using FujiFilm LAS 3000® imager (Fuji, Japan) and their quantification were done with ImageJ. (Antibodies are listed in Supplementary table 1).

### ROS production evaluation

CellROX™ Deep Red Reagent (Thermofisher, Belgium) was added to 3,000 cells seeded in each well of a 96-well plate and treated with APR-246 (25µmol/l), COTI-2 (25nmol/l) and cisplatin (5µmol/l), alone or in combination, each condition having 3 technical replicates. Plates were processed in the Incucyte® S3 Live-Cell Analysis System (Sartorius, Belgium). After a 24-hour incubation, half of plates were irradiated with one fraction of 5Gy, while the other half of plates remained un irradiated. Pictures were taken every 24 hours. Analyses were performed using Incucyte® software.

### GSH measurement

50,000 cells were seeded in each well of the six well plates and were treated with APR-246 (40µM), COTI-2 (400nM) and BSO (40µM) overnight at 37°C, each condition having two technical replicates. Freeze-thaw cycles were used to lyse the cells after which both reduced and oxidized glutathione were measured at 405 nm in a GloMax® spectrophotometer (Promega, Belgium) according to the manufacturer’s instructions (Glutathione Assay Kit, Sigma Aldrich, Belgium).

### Lipid peroxidation evaluation

Lipid peroxidation was assessed using the ImageIT lipid peroxidation kit (Invitrogen, Belgium) containing BODIPY™ C11 as described elsewhere ^22^. 96 well plates containing 4,000 cells/well were exposed to 10µM of Bodipy C11 for 30 minutes to allow its uptake into the cell membrane. Subsequently, the supernatant was removed and medium containing 10µM APR-246, 25µM COTI-2, 3µM cisplatin or 25µM erastin was added, either alone or in combination. Wells with only one effector molecule were tested in three technical replicates while combinations of effectors were tested with six technical replicates. Additionally, one of each pair of treated plates was irradiated with a fraction of 5Gy. The spectrum shift of Bodipy C11 upon peroxidation, from 590nm to 511nm, was recorded using the Incucyte® S3 real time imaging machine. Analysis was performed using the accompanying Incucyte® software.

### Animal experiments

Experimental protocols were conducted in accordance with the European Union Guidelines. All *in vivo* experiments were approved by the local Animal Ethics Committee “Comité d’éthique du Bien-Etre Animal-Université Libre de Bruxelles” (CEBEA) under protocol No 840N. Two million FaDu cells embedded in Cultrex™ matrigel (R&D systems, Belgium) were subcutaneously injected in the hind leg of five to six-week-old female nude (nu/nu) mice (Charles River, France). When tumours reached a volume of about 100mm³, mice were intraperitoneally injected daily with APR-246 (100mg/kg) or COTI-2 (3mg/kg) and weekly with cisplatin (1mg/kg), alone or in combination. Additionally, mice were irradiated with 2Gy for three consecutive days for a total dose of 6 Gy using a SmART^+^ microirradiator (Precision X-Ray Inc., Connecticut, USA) using X-rays at an energy of 225 kV, generated by a current of 13 mA producing a dose rate of 0.3Gy/min at the isocentre. To ensure geometric precision and delivery, each mouse received a position CT scan. The treatment planning software (SmART-ATP, SmART Scientific Solutions, The Netherlands) was used to place the isocentre, to delineate the tumour volume and to do the dosimetry for the irradiation. During the CT scan and irradiation, anaesthesia was maintained using a nose cone (0.5 L/min Oxygen with 1.5% isoflurane). Furthermore, the mouse weight was monitored as well as tumour volume and weight lost.

### Statistics

Statistics were performed using the Graphpad Prism software (MA, USA) in which Student’s t-test was used on Annexin V/7-AAD assay, Glutathione assay, ROS assay, ferroptosis assay; and two ways ANOVA on Clonogenic assays, 3D spheroid culture and *in vivo* assay. Graphs are presented as Mean ± SEM. (* p ≤0,05; ** p≤0,01; *** p≤0,001)

## Results

### APR-246 & COTI-2 synergize with chemoradiotherapy in p53 mutated HNSCC cell lines

To determine optimal concentrations of APR-246, COTI-2 and cisplatin, MTT assays were performed on both Detroit-5462 and FaDu cell lines (Suplementary figure 1). Based on those results, concentrations for clonogenic assay were set at 100µM, 50nM and 0.6µM, respectively. Survival curves showed the association of APR-246 or COTI-2 with cisplatin to significantly increased cell death following radiotherapy compared to each compound alone (Figure 1).

**Figure 1:**
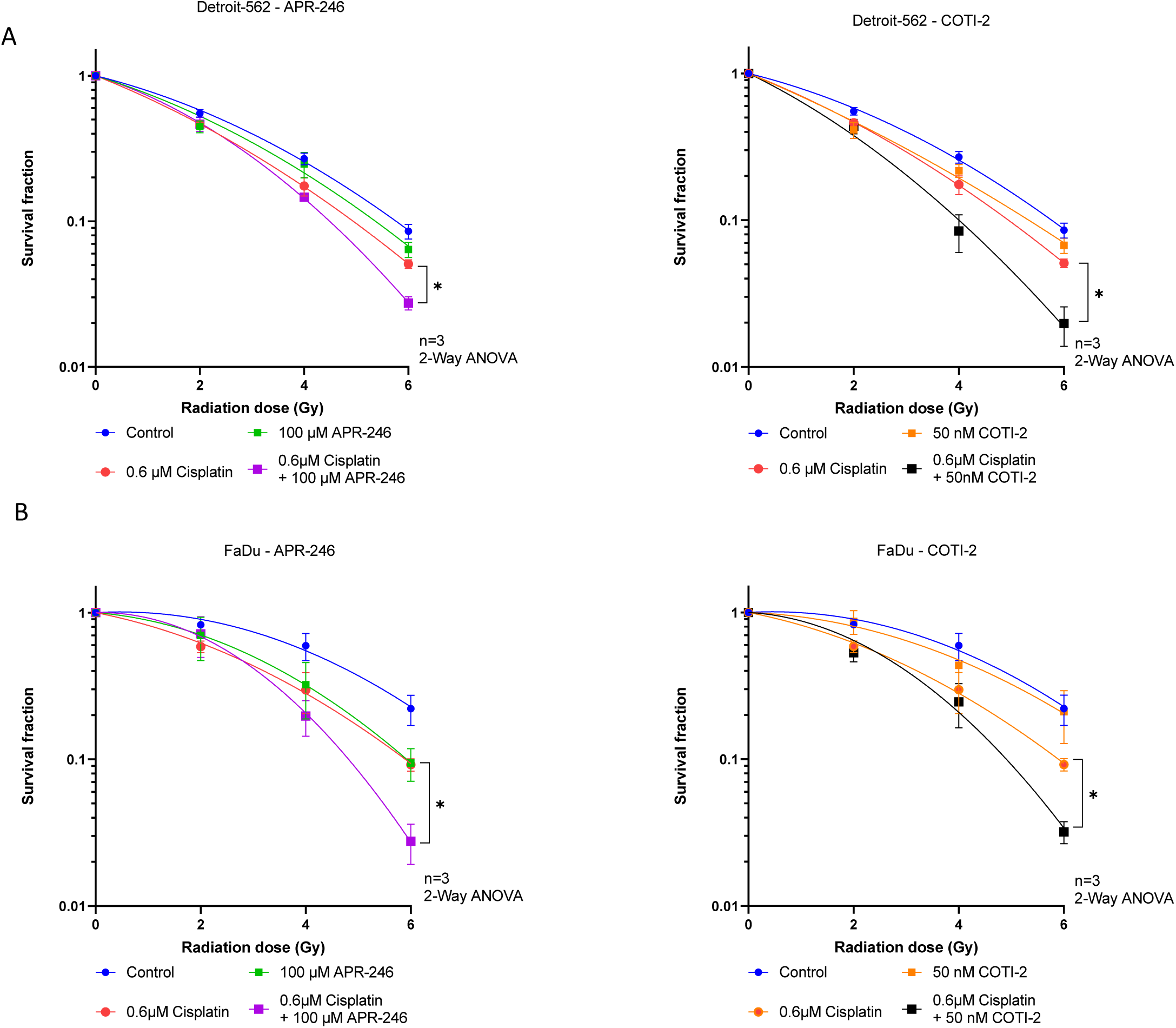
Clonogenic formation of A) FaDu and B) Detroit-562 treated with combinations of Cisplatin and radiotherapy with either Coti-2 or PRIMA-1^Met^ at their respective IC20 concentration with varying doses of radiotherapy. Data is normalized compared to 0Gy and graphs are plotted according to the LQ model. *= p<0,05

Based on the clonogenic results, the CDI (Table 1) and SER at 20 and 50% survival (Table 2 & Supplementary Figure 2 B, D) were calculated. CDI shows COTI-2 to act synergistically with CRT at 4 and 6 Gy in Detroit-562 (0.5± 0.3 & 0.4± 0.3, respectively) and at 6 Gy in FaDU (0.6± 0.1). Similarly, APR-246 synergizes with CRT at 6 Gy (0.6± 0.1) in Detroit-562 and at both 4 and 6 Gy (0.5± 0.2 & 0.2± 0.1 respectively) in FaDu. Additive effects were reported at other doses. This was further supported by SER20 and 50 values. For Detroit-562 treated with APR-246 and CRT the SER20 and 50 are 1.6± 0.1 and 1.8± 0.2, respectively, while for the COTI-2 combined with CRT they are 1.9± 0.2 and 1.9± 0.3 respectively. In FaDu cells, APR-246 with CRT achieved SER 20 and 50 values of 2.6± 0.7 and 4± 2, whereas COTI-2 with CRT reached 2.2± 0.6 and 3± 1, respectively. This shows the COTI-2 CRT combination to be more effective at inducing cell death in Detroit-562 than the APR-246 CRT combination while the opposite is observed in FaDu. Similar results were obtained when calculating the dose required to reach 20 and 50% survival (Supplementary Figure 2 A, C).

**Table 1:** Combination index (CDI) of APR-246 and COTI-2 combined with Chemoradiotherapy in A) Detroit-562 and B) FaDu. Calculated using the survival fraction, n=3 with standard deviation.

| A |  |  | B |  |  |
| --- | --- | --- | --- | --- | --- |
| Dose (Gy) | APR-246 CDI | COTI-2 CDI | Dose (Gy) | APR-246 CDI | COTI-2 CDI |
| 2 | 1.0 ± 0.1 | 0.9 ± 0.1 | 2 | 0.7 ± 0.2 | 0.9 ± 0.3 |
| 4 | 0.9 ± 0.2 | 0.5 ± 0.3 | 4 | 0.5 ± 0.2 | 0.9 ± 0.5 |
| 6 | 0.6 ± 0.1 | 0.4 ± 0.3 | 6 | 0.2 ± 0.1 | 0.4 ± 0.1 |

**Table 2:** Sensitivity enhancement ratio calculated for both A) Detroit-562 and B) FaDu at 20% and 50% survival, n=3 with Standard deviation.

| A |  |  |  |  |  |  |
| --- | --- | --- | --- | --- | --- | --- |
| Detroit-562 | Control | APR-246 | Coti-2 | Cisplatin | Cisplatin + APR-246 | Cisplatin + Coti-2 |
| SER 20% | 1 | 1.08 ± 0.04 | 1.14 ± 0.06 | 1.5 ± 0.2 | 1.6 ± 0.1 | 1.9 ± 0.2 |
| SER 50% | 1 | 1.3 ± 0.3 | 1.4 ± 0.2 | 1.6 ± 0.1 | 1.8 ± 0.2 | 1.9 ± 0.3 |

| B |  |  |  |  |  |  |
| --- | --- | --- | --- | --- | --- | --- |
| FaDu | Control | APR-246 | Coti-2 | Cisplatin | Cisplatin + APR-246 | Cisplatin + Coti-2 |
| SER 20% | 1 | 1.17 ± 0.05 | 1.20 ± 0.05 | 1.3 ± 0.4 | 2.6 ± 0.7 | 2.2 ± 0.6 |
| SER 50% | 1 | 1.3 ± 0.1 | 1.5 ± 0.3 | 1.8 ± 0.7 | 4 ± 2 | 3 ± 1 |

To confirm these sensitizing effects in a 3D model, spheroid cultures were established for both Detroit- 562 and FaDu cell lines transfected with GFP-expressing plasmid. In agreement with previous results, both APR-246 and COTI-2 combined with CRT led to significantly smaller GPF-positive areas in each spheroid image compared to the ones treated by CRT alone, except for FaDu treated with COTI-2 and CRT combination (Figure 2). A two-way ANOVA analysis did not show significant differences between the COTI-2/CRT combination and CRT alone. The combination of COTI-2/CRT only reaches significance when compared with the COTI-2/cisplatin combination (Supplementary table 2).

**Figure 2:**
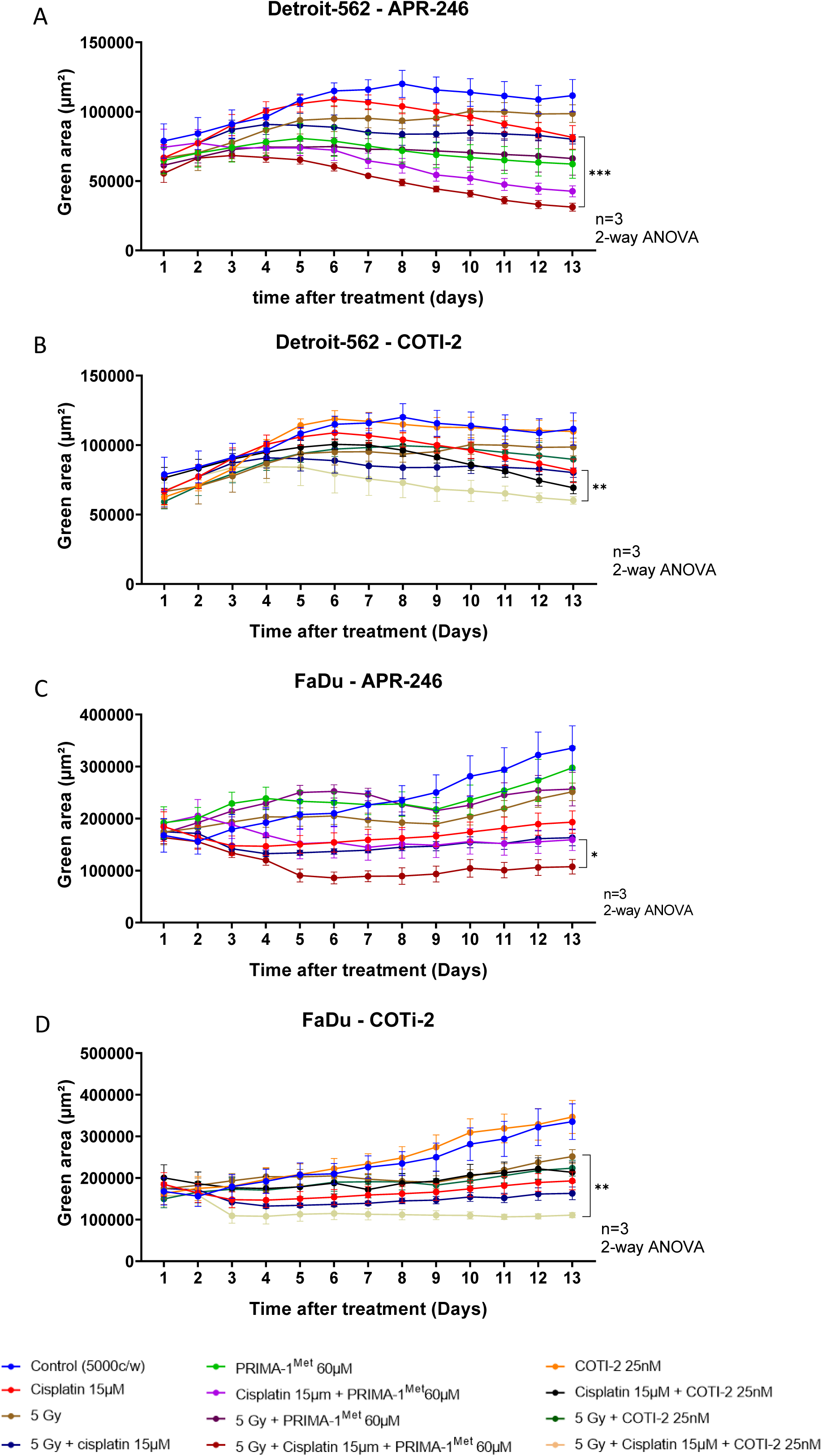
assessment of cytotoxicity in 3D spheroid model formed in ultra low attachement round bottom 96 well plate. The growth was monitored over 13 days. Spheroids created with Dertroit-562 cells treated on day 0 with A) COTI-2 (25nM) or B) PRIMA-1^Met^ (60µM) with cisplatin (15µM) and, on day 1, radiotherapy (5Gy) either alone or in combination. Spheroids made with FaDu cells are treated with C) COTI-2 (25nM) & D) PRIMA-1^Met^ (60µM) with cisplatin (15µM) and radiotherapy (5Gy) either alone or in combination. *=p<0.05 and **=p<0.01.Complete statistical analysis complementary table 1

### APR-246 or COTI-2 in combination with CRT does not induce apoptosis due to a lack of p53 reactivation

As APR-246 and COTI-2 are known p53 reactivators ^17,23^, we hypothesized that their combination with CRT might increase p53-mediated apoptosis. To investigate this, an Annexin V/7-AAD assay was conducted to assess apoptotic cell death (Figure 3). However, the reported differences in cell death do not reach statistical significance compared to the control groups, indicating that cell death from APR- 246/CRT and the COTI-2/CRT combinations likely occurs through mechanisms other than apoptosis.

**Figure 3:**
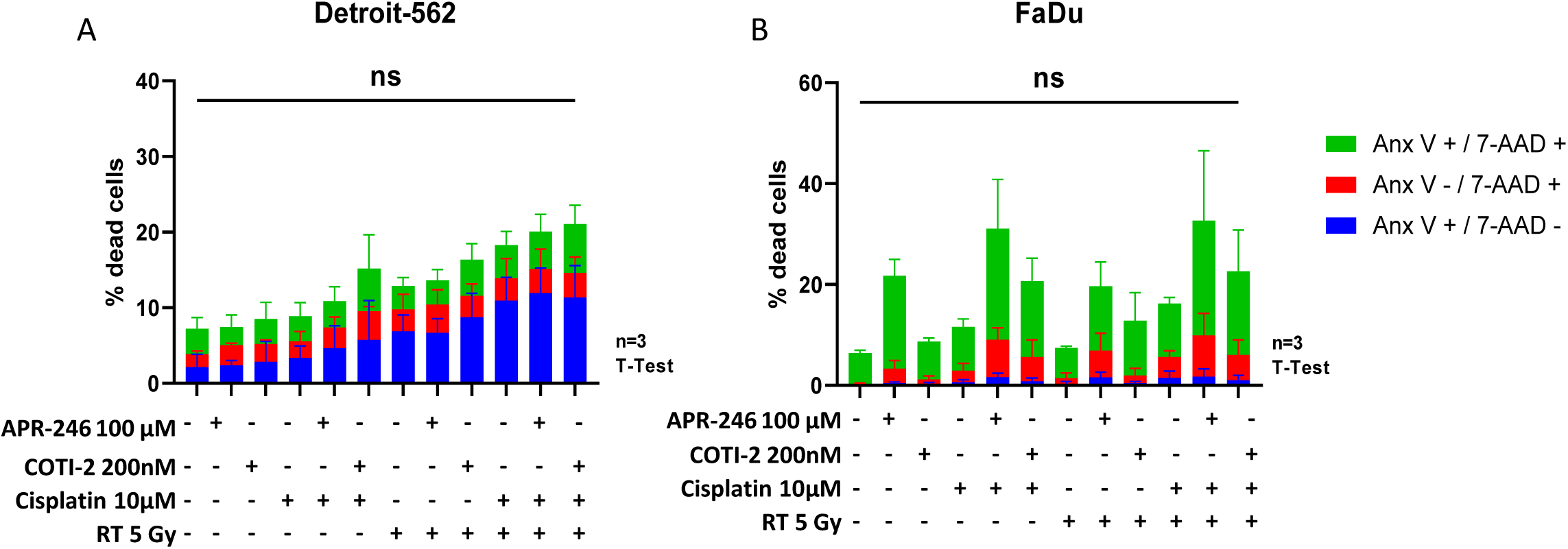
In both A) Detroit-562 and B) FaDu, Neither APR-246 or COTI-2 either alone or in combination with CRT induce significant increases in apoptosis in Detroit-562 obseved via a Annexin V / 7-AAD assay. Statisctical analysis with t-test n=3.

To ensure that treatment with APR-246 or COTI-2 reactivates p53 in Detroit-562 and FaDu, a western blot analysis of p53 and its downstream protein p21 was performed 24 hours after cell treatment with increasing concentrations of APR-246 (25, 50 and 100µM) and COTI-2 (50, 100 and 1000nM). Only Detroit-562 cells treated with 1000nM of COTI-2 expressed higher levels of p21. In FaDu cells, COTI-2 did not induce p21 expression while APR-246 did not lead to an increased expression of p21 in either cell line (Figure 4). These results indicate that in absence of p53 reactivation, apoptotic pathways are not activated and cell death likely occurs through alternative mechanisms^24^.

**Figure 4:**
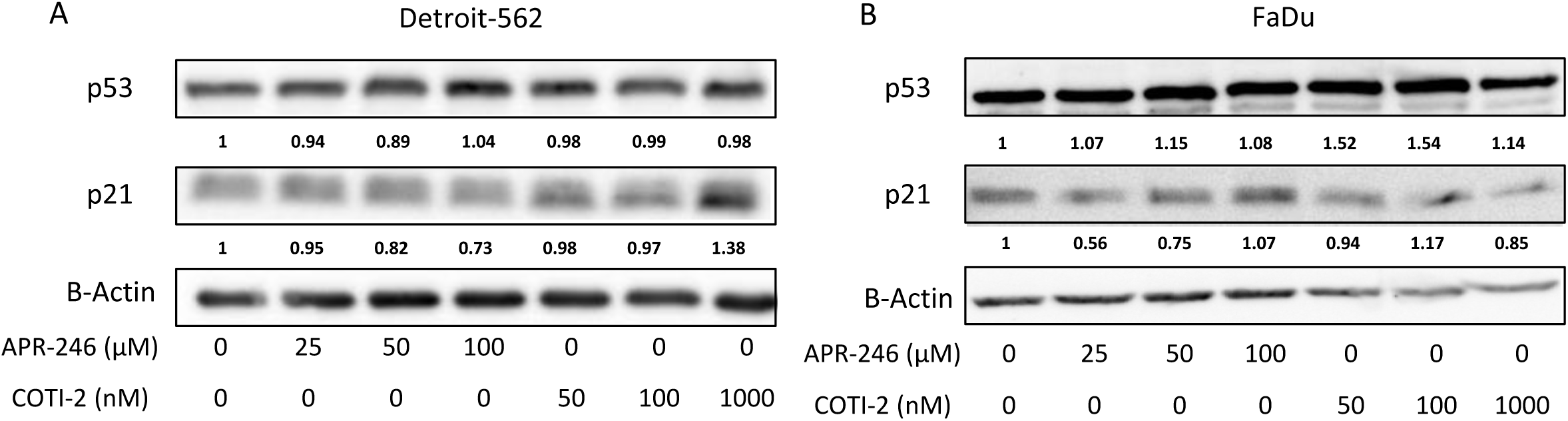
A) Detroit-562 and B) FaDu cells are treated with increasing concentration of APR-246 or COTI-2 24 hours after treatment, cells are lysed and proteins levels are dosed in order to perform p53 and p21 western blots.

### Reactive oxygen species are increased by treatment combination of APR-246 or COTI-2 and CRT irrespective of GSH status

Both radiotherapy and cisplatin are known to induce increased ROS levels. Additionally, APR-246 and COTI-2 have both been shown to influence the redox balance by decreasing GSH levels in the cell. To assess ROS production when combining APR-246 or COTI-2 with CRT, live cell imaging was performed on Detroit-562 and FaDu cells using the CellROX deep red probe (Figure 5). In Detroit-562 cells, ROS levels were significantly elevated with both APR-246 and COTI-2 in combination with CRT, mainly driven by cisplatin (Figure 5A). In FaDu cells, ROS levels seem to be less influenced by these compounds, only reaching significance, for the combinations of APR-246 with CRT compared to CRT alone, at the longest time point post-RT. All monotherapies, except for radiotherapy, we observe significantly increased ROS level compared to the untreated control (Supplementary Table 3).

**Figure 5:**
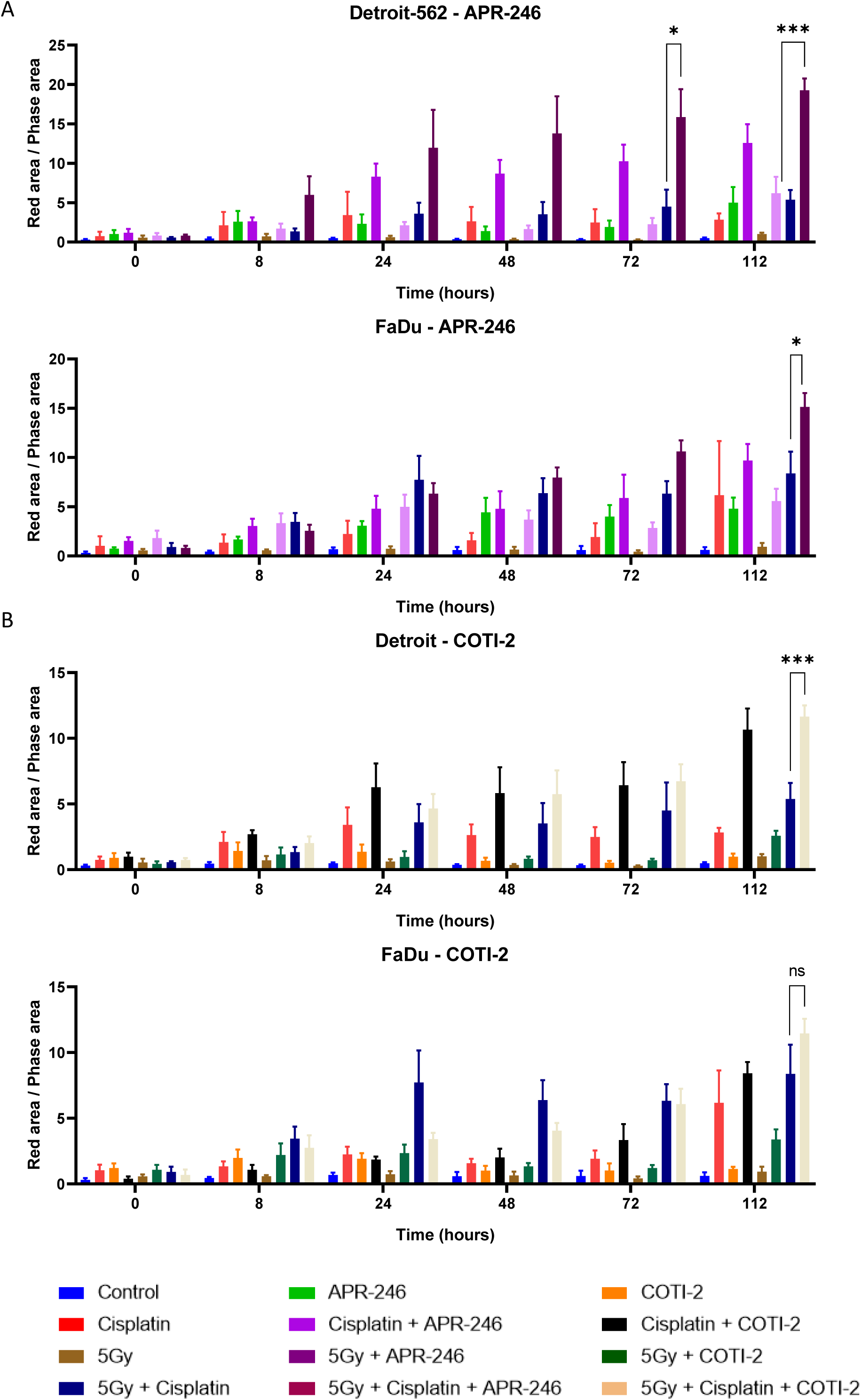
ROS generationa in Detroit-562 and FaDu cells after treatment with A) APR-246 or B) COTI-2 either alone or in combination with cisplatin and radiotherapy. Measurements taken at time after irradiation. Statistics are shown for the triple combination compared to 5Gy + cisplatin.. *=p<0.05, **=p<0.01 or ***=p<0.001. Complete statistics in supplementary table 3

Since increased ROS levels could be an indication of GSH balance disruption, we examined GSH and its oxidized form (GSSG) levels with APR-246 and COTI-2. Detroit-562 cells showed no significant changes in GSH or GSSG levels following APR-246 or COTI-2 treatment (Figure 6A), while FaDu cells exhibited a decrease in GSH without a corresponding increase in GSSG (Figure 6B).

**Figure 6:**
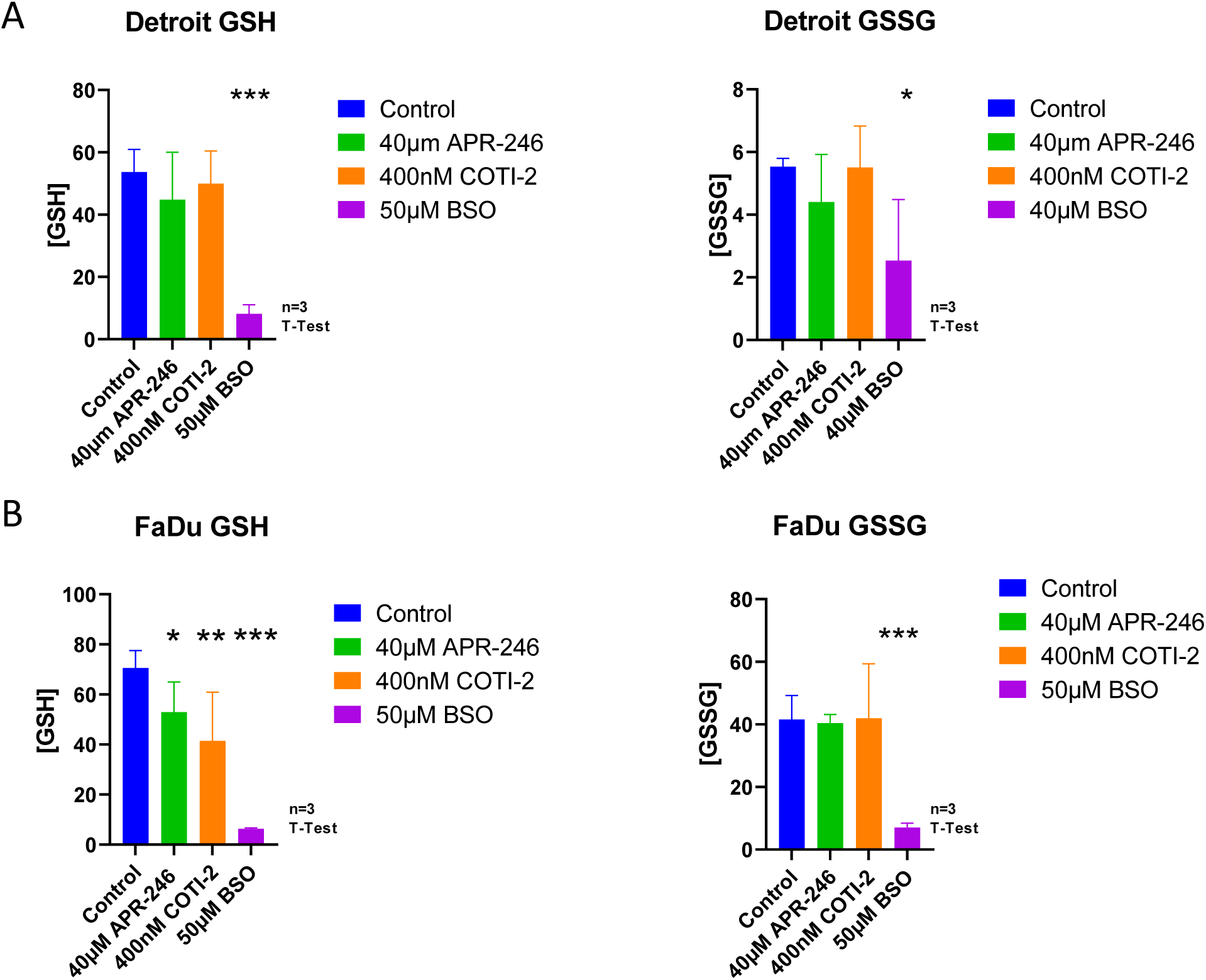
GSH and GSSG are measured in both A) Detroit-562 and B) FaDu cells are treated with elevated concentrations of APR-246, COTI-2 or BSO for 16 hours before lysis and the assay. *=p<0.05, **=p<0.01 or ***=p<0.001.

### Increased ROS and decreased intracellular ferritin suggest potential for ferroptosis in combination treatments

Key markers of ferroptosis, including, proteins involved in intracellular Iron homeostasis (DMT1, FTH1 & NCOA4), cystine/glutamate antiporter (SLC3A2 and xCT complex) and oxidative stress regulators (KEAP1 and NRF2) were investigated following treatment with APR-246 or COTI-2 with CRT.

In Detroit-562 cells treated with APR-246 and CRT, NCOA4 expression decreased by 43% compared to untreated control, while DMT1 and FTH1 remain relatively stable with a 27% and 9% increase in expression respectively. Similarly, SLC3A2 and xCT expression remains stable, showing minor decreases, while KEAP1 is upregulated by 42% with non-significant changes in NRF2 expression. In contrast, when the same cells are treated with COTI-2 and CRT, an increase in FTH1, DMT1 and NCOA4 expression of 32%, 70% and 13%, respectively was observed. This goes with a 21% decrease in SLC3A2 expression and a 34% increase in xCT levels. Like with APR-246, we also reported a 76% increase in KEAP1 levels and a decrease in NRF2 expression by 27% (Figure 7).

**Figure 7:**
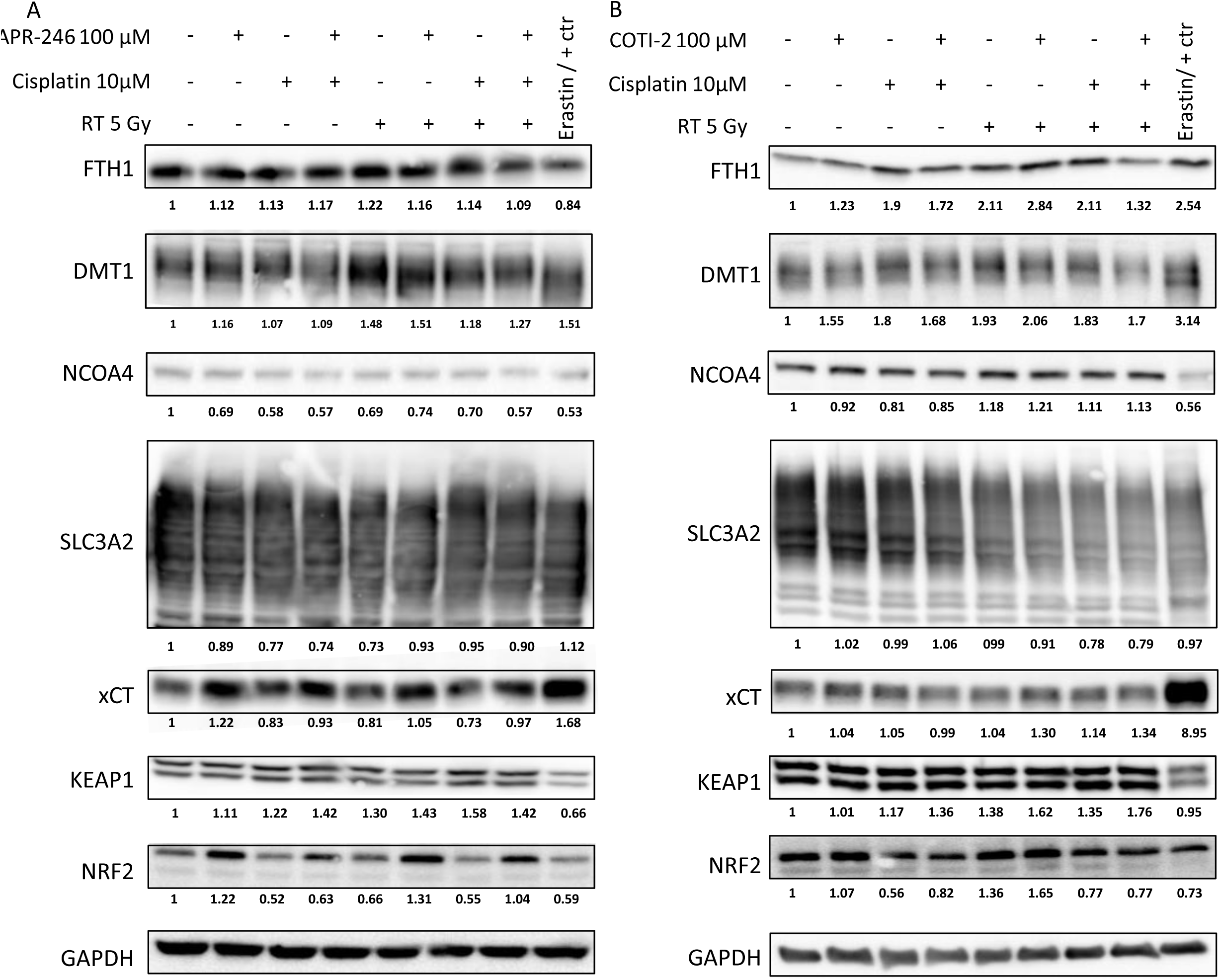
Detroit-562 cells are treated with A) APR-246 or B) COTI-2 either alone or in combination with cisplatin and radiotherapy. Cells lysed 16 hour after irradiation. Markers related to intraellular Iron levels (FTH1, dMT1 & NCOA4), ROS metabolism (KEAP1 & NRF2) and Cyteine import (SLC3A2 &xCT) are detected. Bands are quantified and relativisied compared to B-Actin. Subsequent fold change compared to the control is calculated.

In Fadu cells, both APR-246 and COTI-2 combination treatments result in FTH1 decreases by 76% and 86%, respectively, while a 45% increase in DMT1 expression was reported for the COTI-2 CRT combination treated cells and a 37% decrease in expression for APR-246 CRT combination treated cells. NCOA4 expression decreases by 22% for APR-246 CRT combination treated cells while COTI-2 CRT combination treated cells express 58% more NCOA4, which is responsible for the breakdown of FTH1. However, the cysteine glutamate antiporter complex seems to be affected with a 43% decrease in expression of xCT for COTI-2 CRT combination treated cells and 21% increase in xCT for APR-246 CRT combination treated cells. SLC23A2 expression increases by 77% for APR-246 CRT combination treated cells while COTI-2 CRT combination treated cells maintain stable levels with only a 6% increase in expression. APR-246 CRT combination treatment decreases KEAP1 by 22% while NRF2 expression is increased by 18% for the COTI-2 CRT combination. The COTI-2 CRT combination causes a slight decrease in KEAP1 and stable expression of NRF2 (Figure 8).

**Figure 8:**
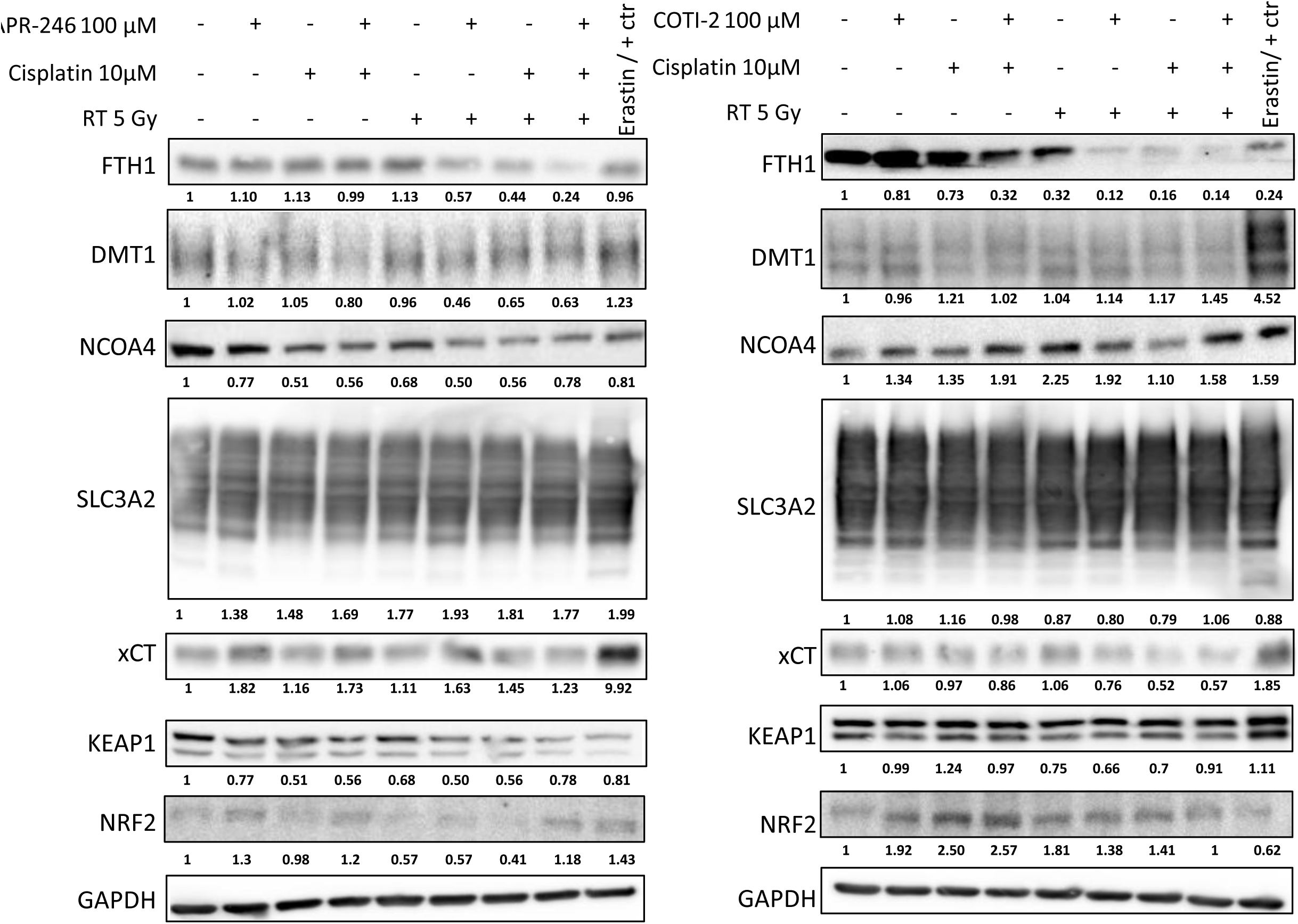
FaDu cells are treated with A) APR-246 or B) COTI-2 either alone or in combination with cisplatin and radiotherapy. Cells lysed 16 hour after irradiation. Markers related to intraellular Iron levels (FTH1, dMT1 & NCOA4), ROS metabolism (KEAP1 & NRF2) and Cyteine import (SLC3A2 &xCT) are detected. Bands are quantified and normalized to their respective GAPDH bands. Subsequent fold change compared to the control is calculated and shown beneath each respective band.

Lipid peroxidation was measured using Bodipy C11, which shifts from green to red as lipids oxidize (Figure 9). Interestingly, both APR-246 and COTI-2 combined with CRT induced higher membrane lipid peroxidation in Detroit-562 and FaDu cells compared to CRT alone.

**Figure 9:**
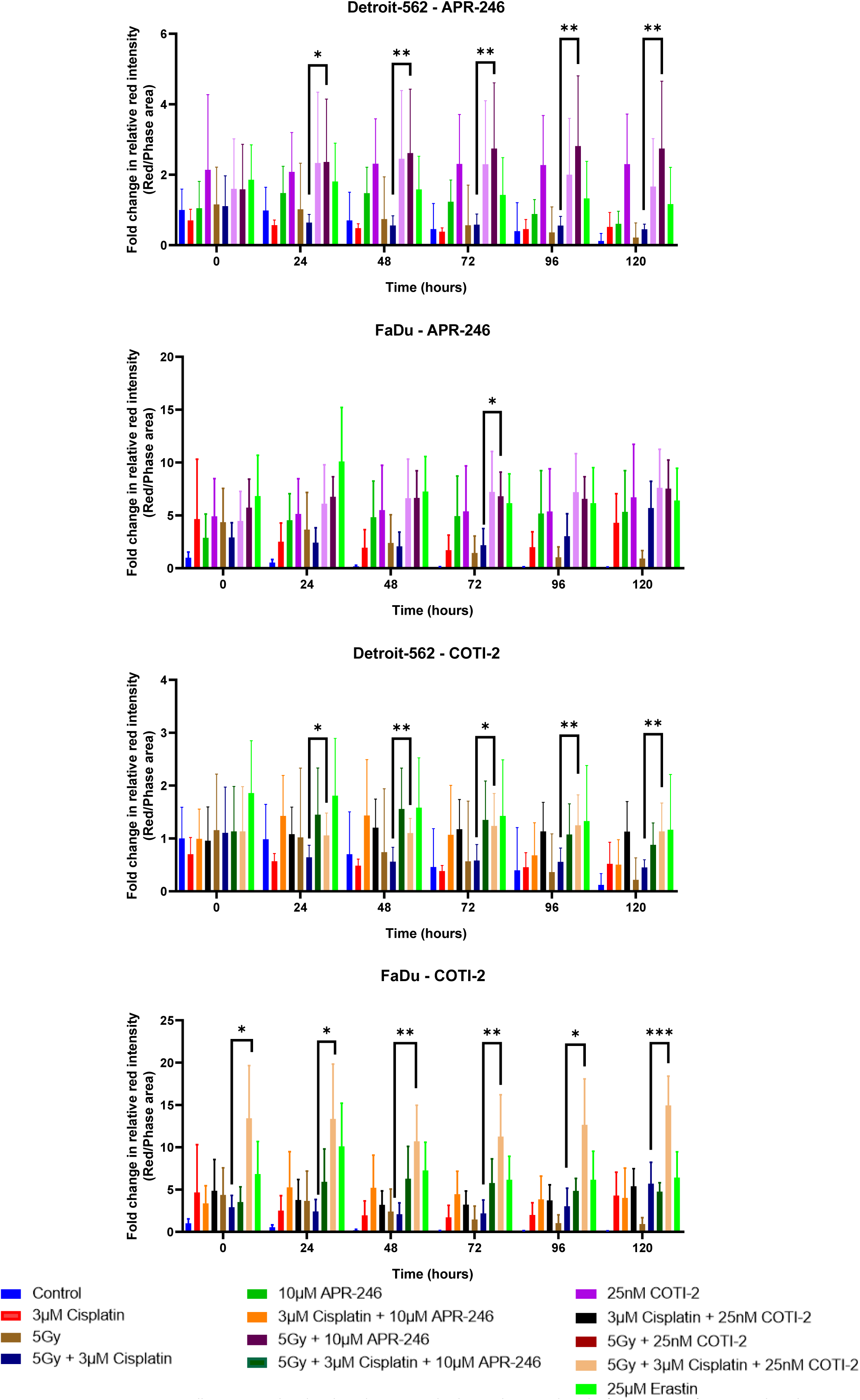
Detroit-562 or FaDu cells are innoculated with Bodipy C11 and subquently treated with A) APR-246 or B) COTI-2 either alone or in combination with cisplatin and radiotherapy. Cells are then monitored using the Incucyte S3 system starting immediately after irradiation; *=p<0.05,

### CRT in combination with APR-246 or COTI-2 increases DNA damage

The combination of APR-246 or COTI-2 with CRT resulted in heightened ROS levels, which, along with cisplatin and radiotherapy’s intrinsic DNA-damaging effects, led to increased DNA damage markers, including γ-H2AX and CHK1 phosphorylation. Western blot analysis performed 1h post-irradiation showed a 2- to 9- fold increase in y-H2AX phosphorylation for both APR-246 CRT and COTI-2 CRT combinations in Detroit-562 and FaDu cells. Similarly,CHK1 phosphorylation increased by 8-30 fold and by 4-10 fold for COTI-2/CRT and APR-246/CRT combinations, respectively (Figure 10).

**Figure 10:**
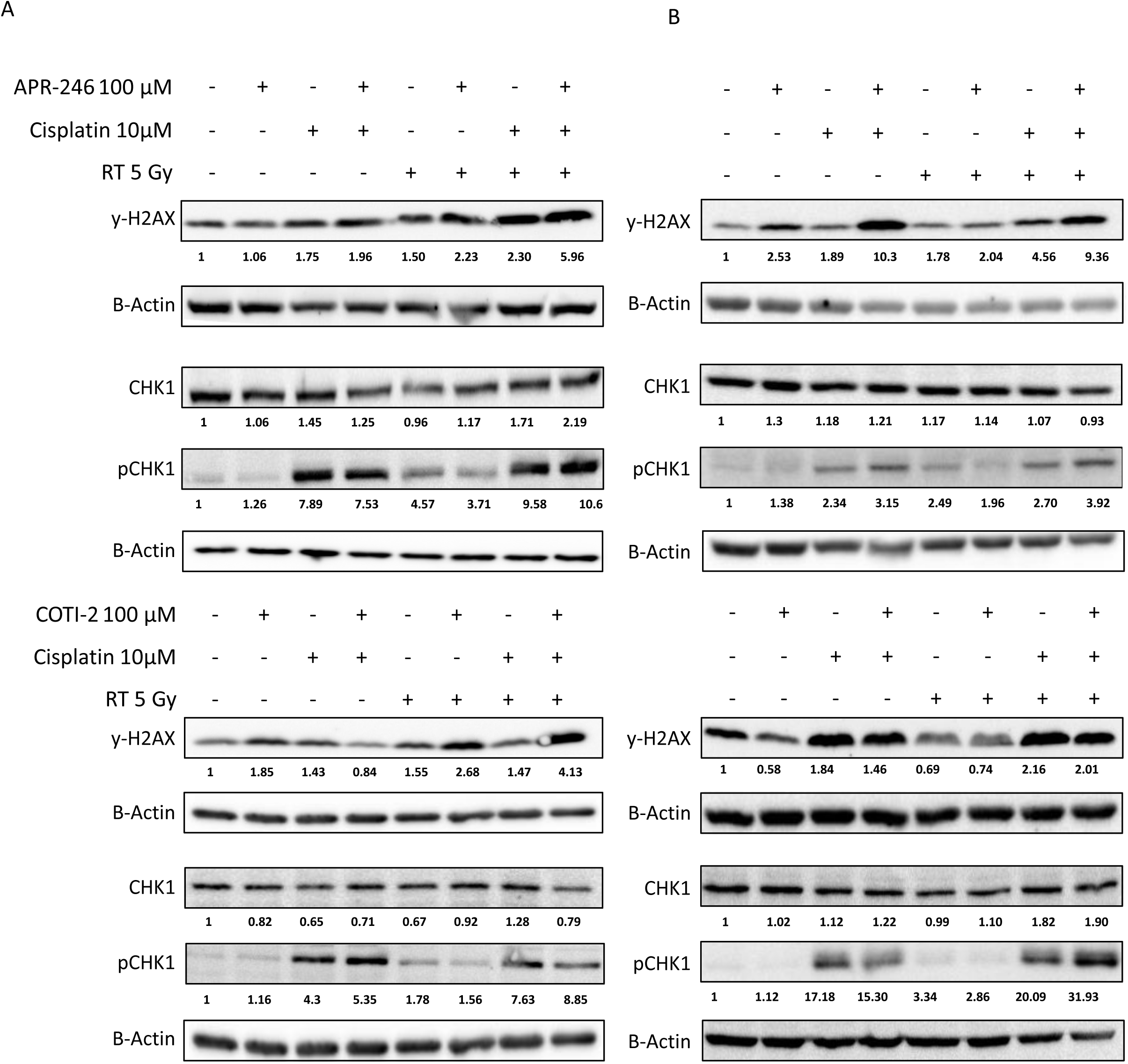
The effect of APR-246 and COTI-2 on DNA damage caused by CRT in both A) Detroit-562 and B) FaDu cells. Cells lysed 1 hour after irradiation. Bands are quantified and relativisied compared to B-Actin. Subsequent fold change compared to the control is calculated.

### APR-246 or COTI-2 with CRT improves tumor control in preclinical murine models

In vivo experiments with FaDu tumor-bearing mice validated the efficacy of APR-246 or COTI-2 combined with CRT Mice treated with these combinations had significantly smaller tumours than the CRT alone group, highlighting the improvement in tumor control (Figure 11A, B). Throughout the study, bodyweight remained stable or increased, and no severe acute toxicity symptoms were observed (Figure 11C, D), indicating a favourable safety profile for these combination treatments.

**Figure 11:**
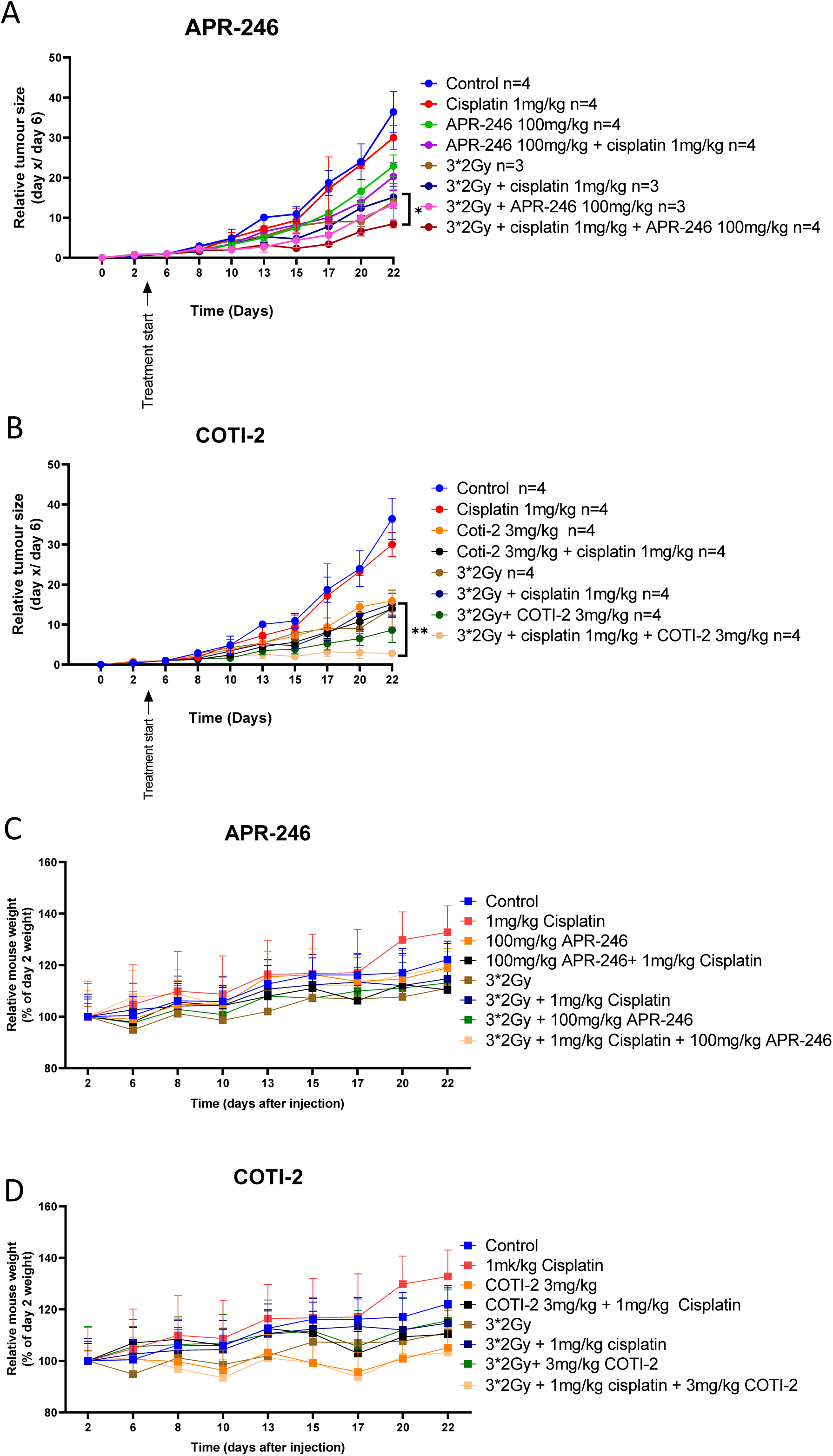
Tumour growth curves of subcutaneously injected HNSCC (FaDu) cells. When tumours reach 100mm³ treatment starts with A) APR-246 and B) COTI-2 with CRT either alone or in combination. Relative mouse weight of mice treated with C) APR-246 or D) COTI-2 alone or in combination. *=p<0.05, **=p<0.01 or ***=p<0.001.

## Discussion

Historically, APR-246 and COTI-2 have been separately evaluated in HNSCC in combination with cisplatin and radiotherapy respectively, yet never in combination with chemoradiotherapy^10,27^. Our study reveals that APR-246 and COTI-2 work synergistically with CRT in both 2D culture of Detroit-562 and FaDu cells as well as in 3D spheroid where significant spheroid size reduction was observed. Interestingly, this was observed even without p53 reactivation in FaDu or in Detroit-562 cells treated with APR-246.

According to literature, the Detroit-562 cell line has a p53 R175H mutation, previously identified as reactivatable by APR-246 and COTI-2^23,28^. However, our results showed that even at the most elevated concentration (100 µM), APR-246 did not seem to reactivate p53, potentially due to alternative targets sequestering MQ (active metabolite of APR-246) and preventing its interaction with p53. FaDu cells, harboring a more complex p53 mutation ( Y126-K132 deletion with R248L substitution), showed no evidence of reactivation by neither APR-246 nor COTI-2. This suggests that the observed synergy with CRT in FADU cells is achieved through a p53-independent mechanism.

Both APR-246 and COTI-2 show potential to induce ferroptosis as monotherapies, though definitive evidence is lacking^29,30^. The increase in ROS with and altered intracellular iron pool (decreased FTH1; elevated DMT1 levels), suggests the initiation of Fenton reaction. In addition to the higher level of peroxidized lipids, these results could trigger the induction of ferroptosis, an iron-dependent form of non-apoptotic cell death. The cystine-glutamate antiporter (xCT comprising SLC3A2 and xCT subunits) mediates cystine import and subsequent GSH biosynthesis, a critical component regulating ferroptosis^31,32^. Although we reported modulation of xCT components by APR-246 and COTI-2 in combination with CRT, increased antiporter activity only occurs when both subunits are upregulated. Finally, KEAP1 upregulation, observed in Detroit-562 cells, inhibits the transcription factor NRF2 and predisposes cells to ferroptosis^33,34^. Overall, combined data from protein expression analyses and lipid peroxidation quantification indicate an enhanced likelihood of ferroptosis triggered by this combination therapy.

Cisplatin and radiotherapy are both known to induce significant amounts of DNA damage which is increased by the addition of both APR-246 and COTI-2. This could be due to the production of radio-induced ROS. As our combination doesn’t induce apoptosis, there is no apoptosis related cleavage of the DNA, therefore, the increased phosphorylation of H2AX and CHK1, which reflect the amount of double and single strand breaks respectively, reflects the increased DNA damage related to increased ROS^36^ If the amount in damage is sufficiently high, it could lead towards increased induction of cell death resulting in the increased cell death observed in both *in vitro* and *in vivo* efficacy experiments.

## Supplementary Figures

**Supplementary Figure 1:**
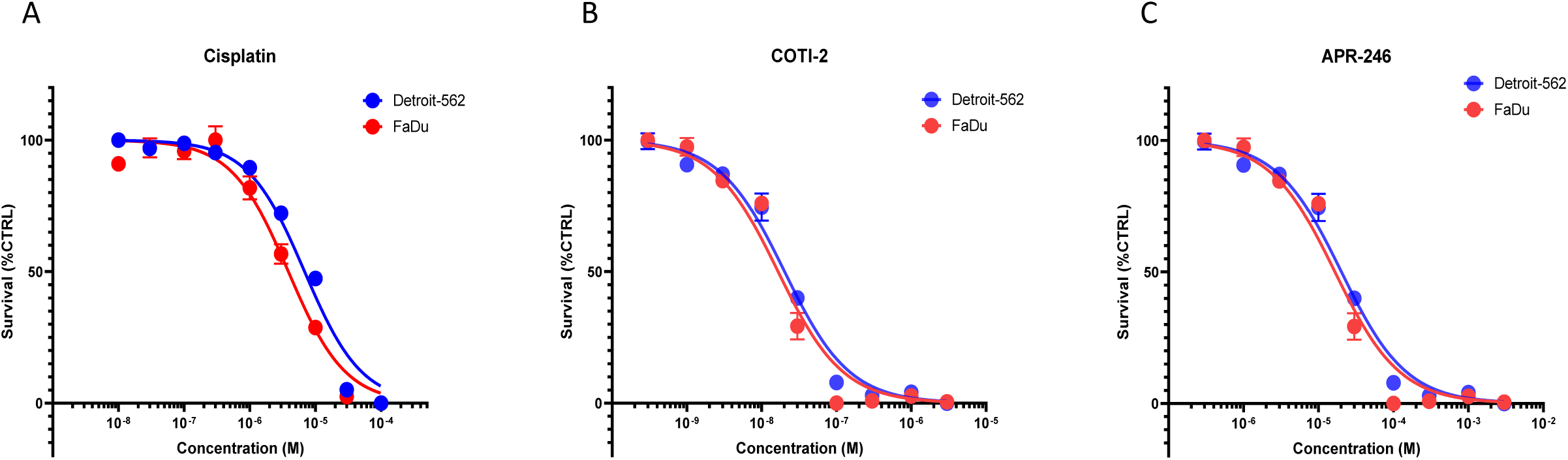
MTT assay for cytotoxic assessment of A) Cisplatin, B) COTI-2 and C) APR-246 on Detroit-562 and FaDu cell lines after 72 hours of treatment.

**Supplementary Figure 2:**
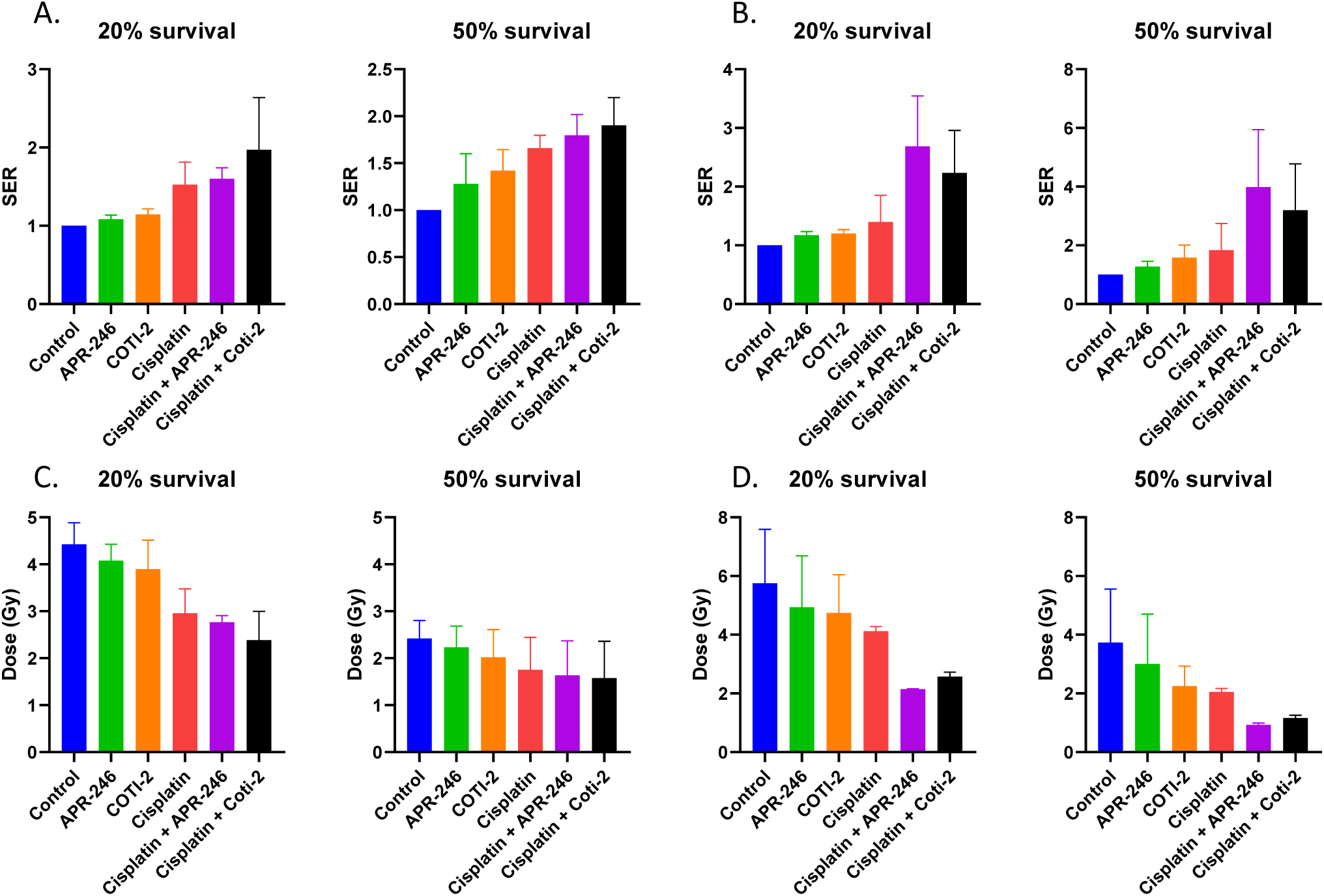
Sensitization enhancement ratio (SER) is calculated by dividing the dose required for 20 and 50% survival by the control in both A) Detroit-562 and B) FaDu. Dose required for 20 and 50% survival for C) Detroit-562 and D) FaDu

**Supplementary Figure 3:**
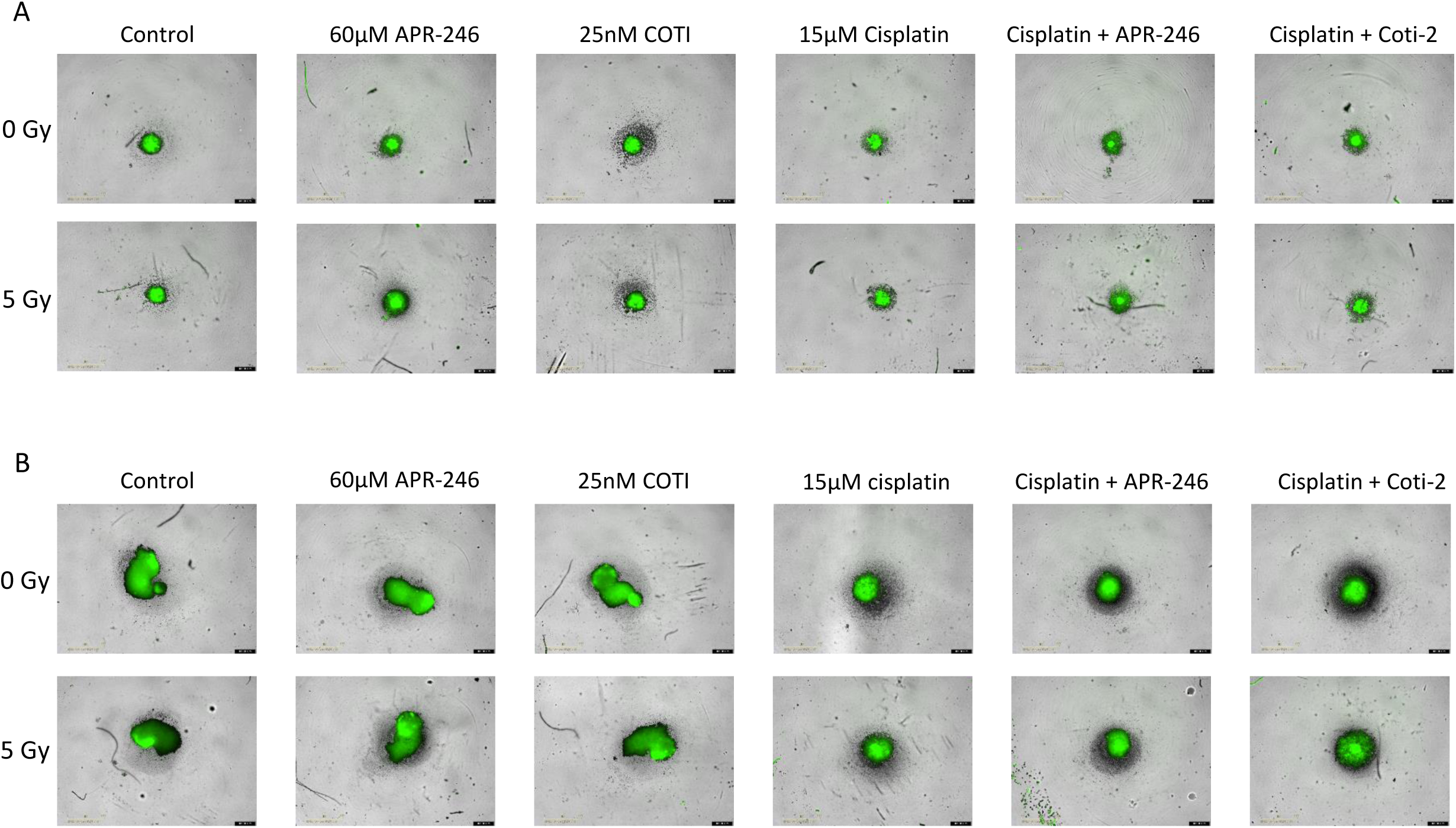
representative images of spheroids in each condition on day 14 for both A) Detroit-562 and B)

**Supplementary table 1:** list of antibodies used in the western blots shown in this paper including their supplier, secondary antibody and dilution used.

| Target | Company | Secondary | Dilution |
| --- | --- | --- | --- |
| P53 | Santa Cruz | Mouse | 1/1000 |
| P21 | Cell signalling | Rabbit | 1/1000 |
| γ-H2AX | Cell signalling | Rabbit | 1/1000 |
| CHK1 | Cell signalling | Rabbit | 1/1000 |
| pCHK1 | Cell signalling | Rabbit | 1/1000 |
| FTH1 | Cell signalling | Rabbit | 1/1000 |
| DMT1 | Cell signalling | Rabbit | 1/1000 |
| NCOA4 | Cell signalling | Rabbit | 1/1000 |
| SLC | Cell signalling | Rabbit | 1/1000 |
| xCT | Cell signalling | Rabbit | 1/1000 |
| KEAP1 | Cell signalling | Rabbit | 1/1000 |
| Nrf2 | Cell signalling | Rabbit | 1/1000 |
| GAPDH | Cell signalling | Rabbit | 1/1000 |
| B-actin | Cell signalling | Rabbit | 1/1000 |

**Supplementary table 2:** Complete statistical analysis of Spheroid assay of both A) Detroit-562 and B) FaDu. P values with p<0.05 are considered significant and shown in green.

| Spheroids day 14 | Control | Cisplatin 15 | APR-246 | coti 25nM | Cisplatin 15 + APR-246 | Cisplatin 15 coti 25nM | RT | RT + Cisplatin 15 | RT + APR-246 | rt + coti 25nM | RT + Cisplatin 15 + APR-246 | rt + Cisplatin 15 + coti 25nM |
| --- | --- | --- | --- | --- | --- | --- | --- | --- | --- | --- | --- | --- |
| Control |  |  |  |  |  |  |  |  |  |  |  |  |
| Cisplatin 15 | 0.6375 |  |  |  |  |  |  |  |  |  |  |  |
| APR-246 | 0.1308 | 0.9342 |  |  |  |  |  |  |  |  |  |  |
| coti 25nM | >0.9999 | 0.4228 | 0.052 |  |  |  |  |  |  |  |  |  |
| Cisplatin 15 + APR-246 | 0.0037 | 0.0624 | 0.7921 | 0.0001 |  |  |  |  |  |  |  |  |
| Cisplatin 15 coti 25nM | 0.1147 | 0.964 | 0.9999 | 0.0128 | 0.015 |  |  |  |  |  |  |  |
| RT | 0.9962 | 0.879 | 0.187 | 0.9895 | <0.0001 | 0.0345 |  |  |  |  |  |  |
| RT + Cisplatin 15 | 0.3745 | >0.9999 | 0.8501 | 0.0936 | 0.0004 | 0.7038 | 0.3667 |  |  |  |  |  |
| RT + APR-246 | 0.2917 | 0.9937 | >0.9999 | 0.1799 | 0.7558 | >0.9999 | 0.4726 | 0.9864 |  |  |  |  |
| rt + coti 25nM | 0.9481 | >0.9999 | 0.7462 | 0.8987 | 0.0284 | 0.7778 | 0.9997 | 0.998 | 0.9271 |  |  |  |
| RT + Cisplatin 15 + APR-246 | 0.001 | 0.0153 | 0.2499 | <0.0001 | 0.5071 | <0.0001 | <0.0001 | <0.0001 | 0.2868 | 0.0052 |  |  |
| rt + Cisplatin 15 + coti 25nM | 0.0332 | 0.5038 | >0.9999 | 0.0023 | 0.1017 | 0.8078 | 0.0013 | 0.01 | >0.9999 | 0.2979 | 0.0001 |  |

**B.**
| Spheroids day 14 | Control | Cisplatin 15 | APR-246 | coti 25nM | Cisplatin 15 + APR-246 | Cisplatin 15 coti 25nM | RT | RT + Cisplatin 15 | RT + APR-246 | rt + coti 25nM | RT + Cisplatin 15 + APR-246 | rt + Cisplatin 15 + coti 25nM |
| --- | --- | --- | --- | --- | --- | --- | --- | --- | --- | --- | --- | --- |
| Control |  |  |  |  |  |  |  |  |  |  |  |  |
| Cisplatin 15 | 0.2015 |  |  |  |  |  |  |  |  |  |  |  |
| APR-246 | >0.9999 | 0.548 |  |  |  |  |  |  |  |  |  |  |
| coti 25nM | >0.9999 | 0.0963 | 0.999 |  |  |  |  |  |  |  |  |  |
| Cisplatin 15 + APR-246 | 0.0584 | 0.9887 | 0.1991 | 0.0234 |  |  |  |  |  |  |  |  |
| Cisplatin 15 coti 25nM | 0.4129 | >0.9999 | 0.8273 | 0.2372 | 0.8493 |  |  |  |  |  |  |  |
| RT | 0.7826 | 0.6076 | 0.9938 | 0.5716 | 0.0842 | 0.975 |  |  |  |  |  |  |
| RT + Cisplatin 15 | 0.0575 | 0.9886 | 0.1926 | 0.023 | >0.9999 | 0.8236 | 0.0356 |  |  |  |  |  |
| RT + APR-246 | 0.9336 | 0.8763 | 0.9996 | 0.8248 | 0.3722 | 0.9937 | >0.9999 | 0.3416 |  |  |  |  |
| rt + coti 25nM | 0.4656 | 0.9937 | 0.8776 | 0.2722 | 0.5182 | >0.9999 | 0.9915 | 0.4161 | 0.9986 |  |  |  |
| RT + Cisplatin 15 + APR-246 | 0.007 | 0.0938 | 0.0231 | 0.0024 | 0.6422 | 0.0419 | 0.0002 | 0.3073 | 0.0213 | 0.0061 |  |  |
| rt + Cisplatin 15 + coti 25nM | 0.0081 | 0.0627 | 0.0249 | 0.0031 | 0.5535 | 0.0323 | 0.0001 | 0.1714 | 0.0214 | 0.0052 | >0.9999 |  |

**Supplementary table 3:**
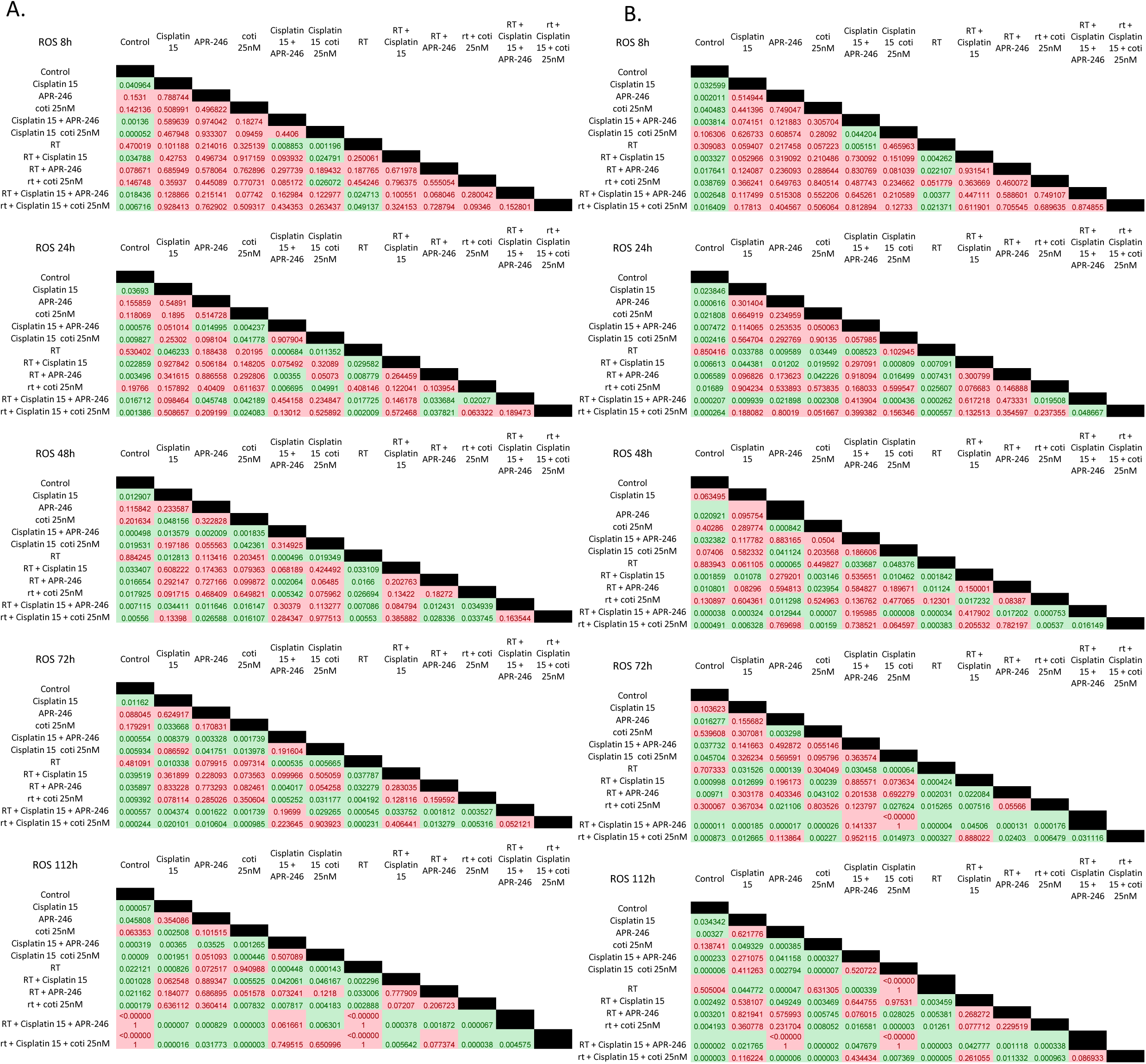
Complete statistical analysis of ROS assay of both A) Detroit-562 and B) FaDu at 8, 24, 48,72 and P values with p<0.05 are considered significant and shown in Green.

**Supplementary table 4:**
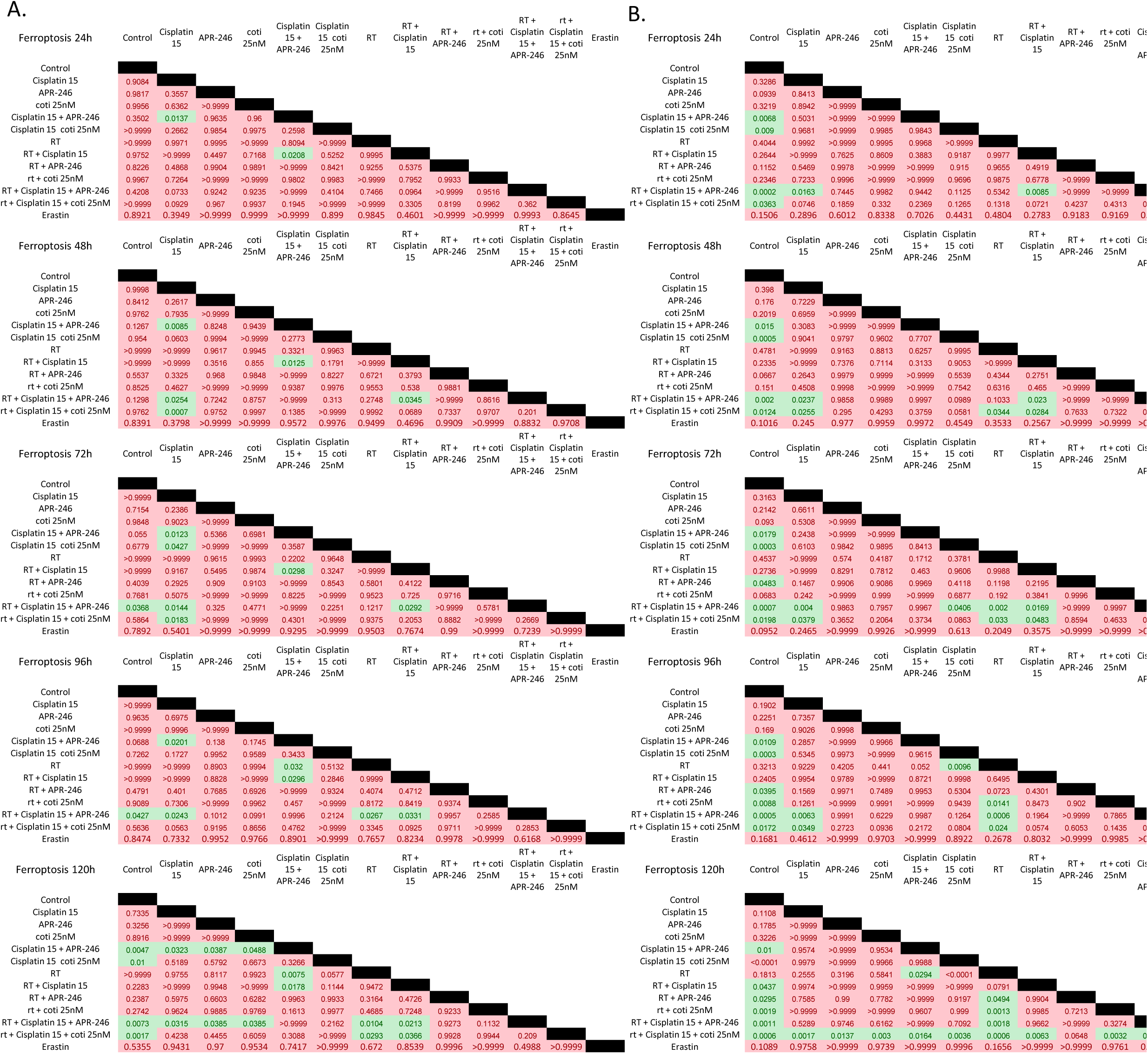
Complete statistical analysis of Ferroptosis assay of both A)Detroit-562 and B) FaDu at24, 48, 72, 9 P values with p<0.05 are considered significant and shown in Green.

